# Cross-species genome-wide identification of evolutionary conserved microProteins

**DOI:** 10.1101/061655

**Authors:** Daniel Straub, Stephan Wenkel

## Abstract

MicroProteins are small single domain proteins that act by engaging their targets into non-productive protein complexes. In order to identify novel microProteins in any sequenced genome of interest, we have developed miPFinder, a program that identifies and classifies potential microProteins. In the past years, several microProteins have been discovered in plants where they are mainly involved in the regulation of development. The miPFinder algorithm identifies all up to date known plant microProteins and extends the microProtein concept to other protein families. Here, we reveal potential microProtein candidates in several plant and animal reference genomes. A large number of these microProteins are species-specific while others evolved early and are evolutionary highly conserved. Most known microProtein genes originated from large ancestral genes by gene duplication, mutation and subsequent degradation. Gene ontology analysis shows that putative microProtein ancestors are often located in the nucleus, and involved in DNA binding and formation of protein complexes. Additionally, microProtein candidates act in plant transcriptional regulation, signal transduction and anatomical structure development. MiPFinder is freely available to find microProteins in any genome and will aid in the identification of novel microProteins in plants and animals

## INTRODUCTION

Genomes of higher eukaryotic organisms encompass on average roughly between 15,000 to 25,000 protein-coding genes. Processes such as alternative splicing, alternative promoter usage, alternative polyadenylation and, at the protein level, proteolytic processing, can significantly increase the number of protein variants these organisms can produce. Furthermore, the formation of higher order protein complexes increases again the functional diversity of proteins. Such higher order protein complexes are often composed of multiple components. Many proteins also associate with different types of complexes in which they 2adopt varying roles. MicroProteins have the ability to interfere with larger proteins and hinder them from engaging in higher order protein complexes, they can also sequester them into other types of complexes. Thus, they are important modulators of biological processes.

MicroProteins exist as individual transcription units in genomes of higher eukaryotes (so called *trans*-microProteins) and most of these transcription units evolved during the evolution of genomes where both whole-genome and local duplications and rearrangements resulted in an amplification of protein-coding sequences followed by a subsequent loss of functional domains (Eguen et al. 2015). In addition, alternative transcription processes such as splicing, promoter choice and 3‘-end processing can also give rise to mRNA isoforms encoding microProteins (so called *cis*-microProteins). In either case, the microProtein is related to a larger protein with different functional domains and interferes with the function of these “precursor proteins” (Eguen et al. 2015).

The first characterized protein that qualifies to be referred to as a microProtein, is the helix-loop-helix (HLH) protein INHIBITOR OF DNA-BINDING (ID). ID was identified almost three decades ago (Benezra et al. 1990) as an interaction partner and inhibitor of basic helix-loop-helix (bHLH) transcription factors. The homotypic interaction of ID with a bHLH transcription factor (through the shared helix-loop-helix domain) renders the latter inactive. The first plant microProteins that were discovered are the LITTLE ZIPPERs (ZPR) proteins, which are small proteins containing a leucine-zipper domain (Wenkel et al. 2007; Kim et al. 2008). ZPR microProteins interact with the much larger class III homeodomain leucine-zipper (HD-ZIPIII) proteins through their leucine-zipper domain and the resulting HD-ZIPIII/ZPR heterodimer is unable to interact with DNA, thus mimicking the ID/bHLH module. In the past years many more microProteins targeting transcription factors have been identified in plants (Eguen et al. 2015). Furthermore, it is possible to design synthetic microProteins that inhibit proteins of interest (Seo et al. 2012). Taken together, these findings indicate that microProtein interference is a powerful way to regulate or fine-tune protein activity.

It seems implausible that microProteins are more abundant in plant genomes when compared to animal genomes or that they exclusively target transcription factors. To identify a larger variety of potential microProteins and microProtein regulatory modules in plant and animal genomes, we have performed a computational approach taking protein size, domain organization and evolutionary origin into account. This approach yielded the identification of 34,840 individual microProtein candidates belonging to 15,867 protein families, with 2,841 in human, 1209 in mouse, 907 in zebrafish, 567 in fruit flies, 1324 in C. elegans, and 1,589 in *Arabidopsis*, 4160 in tomato, 6215 in potato, 1990 in Sorghum, 3447 in rice and 10591 in maize (Suppl. Table 1). This new microProtein dataset provides a valuable resource for investigating mechanisms of microProtein functions in plants and animals and the miPFinder program can be used to analyze new genomes as soon they become available.

## RESULTS

### Key features of microProtein candidates

All microProteins known to date are small in size, ranging from 7 to 17 kDa, overall comprising less than 120 amino acids (Eguen et al. 2015). To exert their function, microProteins require only a single functional domain that acts as a protein-interaction platform to sequester their targets. While the sizes of protein domains vary tremendously, the average maximum length of a protein-interaction domain is approximately 100 amino acids (Wheelan et al. 2000). Considering these values and the fact that all known microProteins are less than 120 amino acid in length, we decided to use a maximum length of 140 amino acids to predict novel microProteins.

A second parameter to take into account when trying to identify novel microProtein candidates is the protein organization of potential targets or ancestor. As described above,trans-microProteins exists as individual transcription units allowing their evolutionary origin be traced back. A good example are the plant-specific ZPR proteins that originate from a large homeodomain leucine-zipper ancestor molecule, which got sequentially shortened by gene duplication, degeneration, and truncation (Floyd et al. 2014). The ZPR-ancestor protein is a multi-domain protein that has the ability to homodimerize. In order to predict potential microProteins, we reasoned that a putative microProtein ancestor protein should be large enough to harbor at least two functional domains, consequently we set a minimum ancestor protein size of 250 amino acids. This step also eliminates the identification of small proteins that belong to protein families in which some members are only marginally larger. Finally, we discovered that searches made with a consensus sequence of related microProtein candidates rather than individual protein sequences against a database of larger proteins significantly increases the sensitivity for identifying distantly related sequences wherefore the microProtein-finder program starts with extracting consensus protein sequences from all small protein families.

### Computational prediction of small and related proteins

In the first step, miPFinder assigns protein sequences as putative microProteins and putative ancestors solely by size. Therefore, the sequence database is divided into small (≤140aa) and large (≥250aa) sequences. Next, BLAST searches with all small sequences against each other are being performed, resulting in the division of microProteins in single-copy proteins and groups of related sequences (BLAST, cutoff e-value ≤0.001). Each group of small proteins is subsequently aligned (clustalw, gap opening penalty = 20, no end gap separation penalty), combined to a consensus profile (hmmbuild) and compared to all large proteins (hmmsearch, cutoff e-value ≤0.1 and c-Evalue ≤0.05). For un-grouped small sequences (single copy microProteins), similar large proteins are chosen based on the initial BLAST search. Grouped or un-grouped small sequences are considered “microProtein candidate families” and included in further analysis only if they are similar to at least one larger putative ancestor. All putative ancestors are reported in order of significance and up to ten putative ancestors and their microProtein candidate family are realigned (clustalw, gap 5 opening penalty = 20), rated, and linked in the final report. Additionally, the e-value of the microProtein-ancestor search is stated, which might help in the manual evaluation of microProtein candidates when prioritizing on highly significant similarities.

In addition to the significance values (BLASTP/hmmsearch e-value), we created a rating system that favors known microProteins. This rating is based on the clustalw alignment of the microProtein candidate family and their putative ancestor(s). First, conserved regions (small proteins and ≤10 similar segments of large proteins, blastp/hmmsearch) are aligned (clustalw) and regions with low gap content (length ≥20 aa and gaps ≤10%) are extracted. This step enriches for regions with high similarity and extracts potential domains. Next, each microProtein candidates and putative ancestors are extracted and two consensus sequences are assembled. The similarity of the consensus sequences is rated based on the Blosum62 table and the following equation:

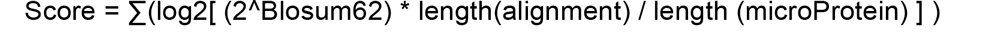

Here, the score is modified by the alignment length in proportion to the length of the microProtein candidate. The resulting alignment rating favors known microProteins and is inversely related to the e-values (Suppl. Fig 1), that is a low e-value corresponds to a high microProtein alignment rating.

### MicroProteins with protein-interaction domains

The core mechanism of microProtein function relies on their ability to interact with respective target proteins. MicroProtein-candidates containing known protein-protein interaction domains, or sequences related to PPI-domains are therefore more likely to function as microProteins compared to small proteins not containing such domains. To identify and annotate protein-protein-interaction domains within microProteins and ancestral proteins, miPFinder utilizes the Pfam and iPfam databases.

MiPFinder assigns Pfam domains to all large proteins (hmmscan, cutoff e-value ≤0.1 and c-Evalue ≤0.05), reports domains that have similarity to microProtein candidates (≥60% length of the Pfam domain) in order of significance, and matches these to interchain interaction domains in iPfam. Domains with interchain interaction properties mediate interactions between amino acid chains, a prerequisite for microProtein function.

### Evolutionary conserved microProteins

Proteins that are conserved in several related species are more likely to have retained a function under evolutionary pressure. Additionally, conserved sequences are less prone to be annotation artifacts or degenerated pseudo-genes. For these reasons, we favor the identification of evolutionary conserved microProteins thus reducing the number of false-positive hits. However, it is important to note, that species-specific microProteins should not be ignored because they could be involved in species-specific traits and in some cases might even have acted as facilitators of speciation.

We employed OrthoFinder (Emms and Kelly 2015) to uncover homology relationships of microProtein candidates among species. OrthoFinder solves biases in whole genome comparisons and is more accurate than other orthogroup inference methods. Like other algorithms it performs sequence comparisons via BLAST but additionally normalizes for gene length and phylogenetic distance in cross species comparisons. OrthoFinder outperforms all other commonly used orthogroup inference methods.

### Detection of known microProteins using miPFinder

All 22 known Arabidopsis microProteins are present in the list of microProtein candidates generated by miPFinder (Table 3). Additionally, ZPR (Wenkel et al. 2007), MIF (Hu and Ma 2006) and MIP1A/MIP1B (Graeff et al. 2016) are exclusively grouped according to their microProtein family associations. However, MYB-microProteins (Tominaga-Wada et al. 2011) and HLH-microProteins (Wang et al. 2009; Zhang et al. 2009) families harbor additional members that have not been studied to date. The latter microProtein family is also misleadingly divided into two groups, however, the group comprises of 10 known microProteins from two clades, the KDR-ILI1-like and the PAR-like subgroup. MYB-microProteins, HLH-microProtein, and MIP1A/MIP1B are correctly reported as being similar to an interaction domain, whereas ZPR (bZIP-TF) and MIF (ZF-HD) domains are not described as interaction domains in iPfam. MiPFinder retains all these microProteins irrespective of their domain but reports the domain and its iPfam annotation. Therefore the researcher can decide whether to rely on iPfam interaction domain annotations or not. ZPR, MIF, Myb-microProteins, and HLH-microProtein (KDR-ILI1-like subgroup) are identified in all plants, while HLH-microProtein (PAR subgroup) and MIP1A/MIP1B only in dicotyledonous plants.

MicroProteins are also known in animals. The first identified microProtein, Inhibitor of DNA binding (ID), was initially identified in mice (Benezra et al. 1990) and miPFinder is able to identify ID2 and ID3 in mouse, however ID1 and ID4 are omitted, because they are too large (148aa and 161aa) (Table 3). ID-like microProteins are properly reported to contain a HLH protein-protein interaction domain. Related microProtein candidates are also found in human and zebrafish.

MicroProteins are not always encoded independently from their large ancestor, *cis*-microProteins are splice variants of a bigger protein that negatively regulate their target. The human cis-microProtein of Regulator of G-protein signaling 5 (RGS5), a small splice variant that can negatively inhibit its targets function (Liang et al. 2005), is not identified, because the supposedly large ancestor RGS5 is shorter (201aa) than miPFinder standard setting allows (≥250aa). To allow for adjustments in microProtein candidate detection, the parameters for the maximum microProtein and minimum ancestor length are easily tunable in miPFinder.

### Identifying novel microProtein candidates with miPFinder

In most protein databases, sequences are derived from translated RNA transcripts, which in some cases represent only truncated versions of full-length mRNA sequences. For human and mouse, proteins encoded by the representative protein-coding “GENCODE Basic” transcript set were used. GENCODE combines manual and automatic annotation and aims to annotate all evidence-based gene features in human and mouse genomes at a high accuracy. GENCODE’s Transcript Support Level highlights the well-supported and poorly supported transcript models and transcripts without any transcriptional evidence were omitted. To deplete incomplete sequences for other organisms, only peptides which were derived from protein-coding nucleotide sequences that contain a start codon (ATG), stop codon (TAA, TGA, TAG), and a length that is a multiple of three were considered.

Incomplete protein sequences were filtered by enriching complete coding sequences. The percentage of transcripts that passed the quality filter varied considerably. In most organisms, more than 98% protein sequences appeared to be complete, however in maize and zebrafish only 91% and 72% of the protein sequences passed the filter. Additionally 60% of human and 72% of mouse transcripts and their corresponding proteins are in Ensembls GENCODE basic set, and of these, approximately 80% are either with transcriptional evidence or not tested for expression (Table 1).

**Table 1.** Overview of miPFinder results.

| Species | Protein sequences |  |  | microProtein candidate families |  |  |  |  |  |  |
| --- | --- | --- | --- | --- | --- | --- | --- | --- | --- | --- |
|  | Original | Filtered<sup>a</sup> | % | Total | Trans-<br>miP<sup>b</sup> | %<sup>c</sup> | PPID<sup>d</sup> | %<sup>c</sup> | ≥50%<br>≥250aa<sup>e</sup> | %<sup>c</sup> |
| *Arabidopsis thaliana* | 35386 | 35364 | 99.94 | 551 | 531 | 96 | 193 | 35 | 328 | 60 |
| *Solanum lycopersicon* | 34727 | 34415 | 99.10 | 1767 | 1767 | 100 | 419 | 24 | 1344 | 76 |
| *Solanum tuberosum* | 51472 | 50631 | 98.37 | 2772 | 1422 | 51 | 587 | 21 | 2011 | 73 |
| *Sorghum bicolor* | 47205 | 46544 | 98.60 | 866 | 861 | 99 | 206 | 24 | 557 | 64 |
| *Oryza sativa* | 52424 | 52417 | 99.99 | 1661 | 1578 | 95 | 305 | 18 | 1090 | 66 |
| *Zea mays* | 88760 | 80694 | 90.91 | 5132 | 3673 | 72 | 1007 | 20 | 3688 | 72 |
| *Homo sapiens* | 101933 | 48542 | 47.62 | 1235 | 320 | 26 | 340 | 28 | 850 | 69 |
| *Mus musculus* | 56337 | 31983 | 56.77 | 526 | 221 | 42 | 186 | 35 | 346 | 66 |
| *Dario rerio* | 44487 | 32031 | 72.00 | 371 | 201 | 54 | 165 | 44 | 253 | 68 |
| *Drosophila*<br>*melanogaster* | 30362 | 30152 | 99.31 | 218 | 128 | 59 | 74 | 34 | 124 | 57 |
| *Caenorhapditis elegans* | 30939 | 30925 | 99.95 | 768 | 372 | 48 | 168 | 22 | 551 | 72 |

<sup>a</sup>coding sequence length is a multiple of 3 and with start and stop codon; for *H. sapiens* and *M. Musculus* protein coding sequences of GENCODE basic that are not flagged as without any transcription evidence.
<sup>b</sup>microProtein candidates do not contain a cis-miP.
<sup>c</sup>percentage of total microProtein candidate families (column „Total“).
<sup>d</sup>sequences with annotated protein-protein interaction domain (PPID).
<sup>e</sup>more than or equal to 50% of related sequences (blast or hmmsearch) are ≥250 aa in length.

In Arabidopsis, miPFinder identified 1,589 microProtein candidates belonging to 195 groups and 356 single small sequences, resulting in a total of 551 microProtein candidate families. Around 34% of these microProtein candidate families exhibit similarities to known protein-protein interaction domains that are related to putative ancestral proteins. In all of the 11 proteomes that we have investigated here, the majority (~67%) of the microProtein candidate families are related to a higher number of large proteins (≥50% ≥250aa in length) than small proteins. In plants, groups without *cis*-microProtein candidates which are alternative products of their ancestor genes, make up the majority of microProteins identified in these species, although in potato and maize these numbers are lower (51% and 72% respectively, see Table 1). In metazoans, small splice variants of large proteins are present in more than half of the microProtein candidate families. For example, only 26% of human candidate microProtein families are exclusively *trans-microProteins.* The number of splice variants per gene, which is significantly higher in mammals than in plants, might explain these differences (Kim et al. 2007). However, invertebrates and plants have a similar proportion of spliced genes (Kim et al. 2007), therefore the difference in this situation might be due to the dissimilar annotation degree of splice variants among the databases.

In contrast to OrthoFinder (Emms and Kelly 2015), which calculates orthogroups with both orthologues and paralogues of all given species, we opted in miPFinder for group formation with relatively relaxed cutoffs. For conservation, individual microProtein candidates were combined with OrthoFinder results. Individual microProtein candidates were sequentially tested for their presence in all 11 species. In total seven microProtein candidate families are conserved in all 11 data sets (Fig 2, red; Table 2). These proteins are involved in basic pathways and have domains resembling RNA recognition motif, Cytochrome b5-like Heme/Steroid binding domain, ubiquitin-like or transcription factor-like domains. However, further investigations showed only limited probability for microProtein function among these groups. For example in most species Cytochrome b5-like Heme/Steroid binding domain and Transcription factor S-II (TFIIS) microProtein candidates show relatively high percentage of small proteins (>66%) and weak similarity to their putative ancestors. Due to evolutionary origin, we expect that microProteins make up only a fraction of large families of proteins. In other words, in a given set of related protein sequences, the number of microProteins should be small, especially in relation to the number of ancestors, but also compared to middle sized proteins. Additionally, sequence similarity to the putative ancestors should be high. The microProtein candidate families that are conserved in all 11 species do not conform to these criteria.

**Figure 1.**
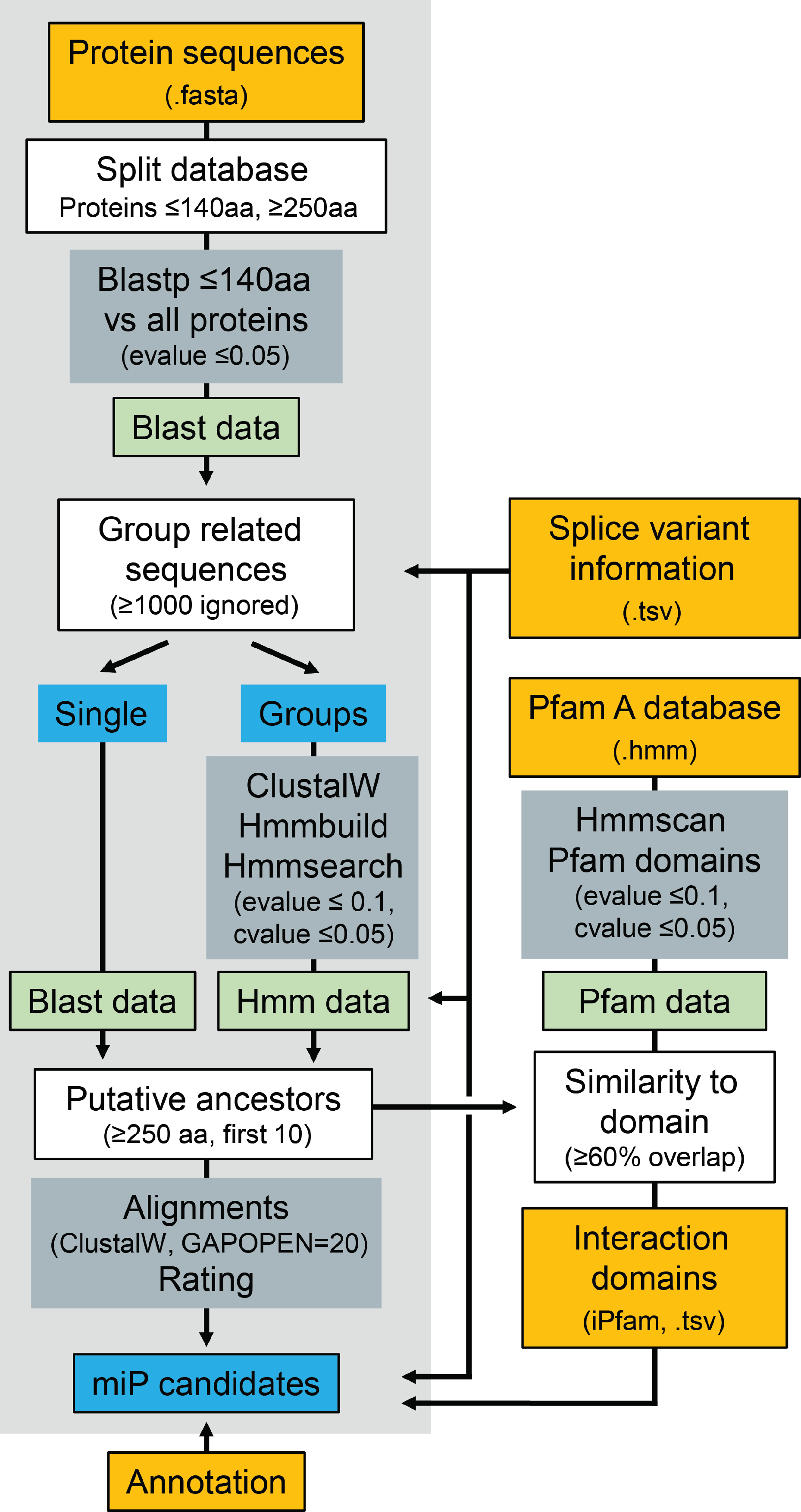
Flow chart miPFinder. Mandatory steps are with a light grey background. Orange: databases, Green: data packages, Grey: tools, Blue: Lists, White: Custom functions.

**Figure 2.**
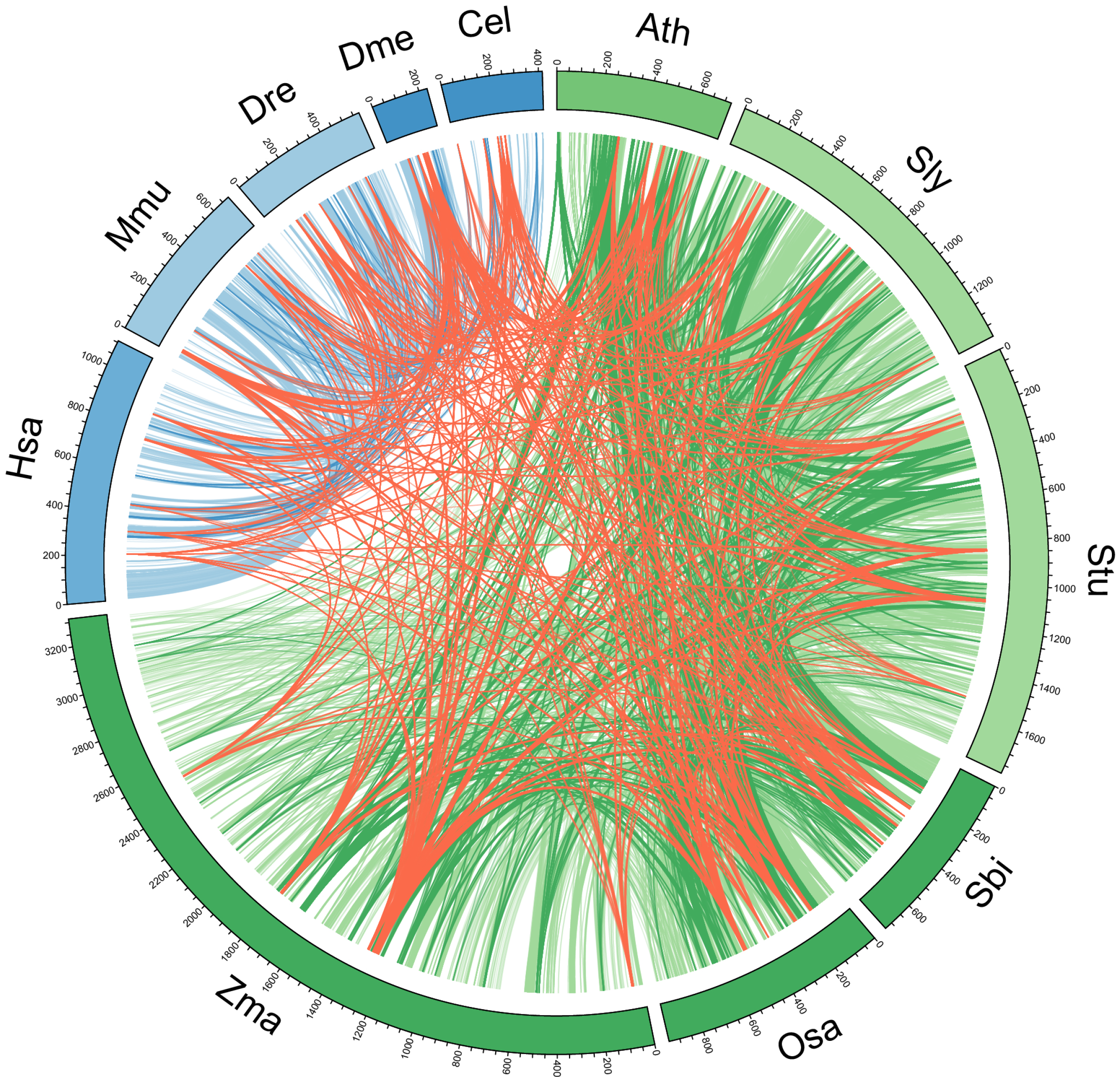
Circos plot of individual microProtein candidates. Links indicate conservation between species based on OrthoFinder. Red: in all 11 species; Dark blue: exclusively in all five metazoans; Light blue: only in metazoans; Dark green: exclusively in all six plants; Light green: only in plants.

**Table 2.**
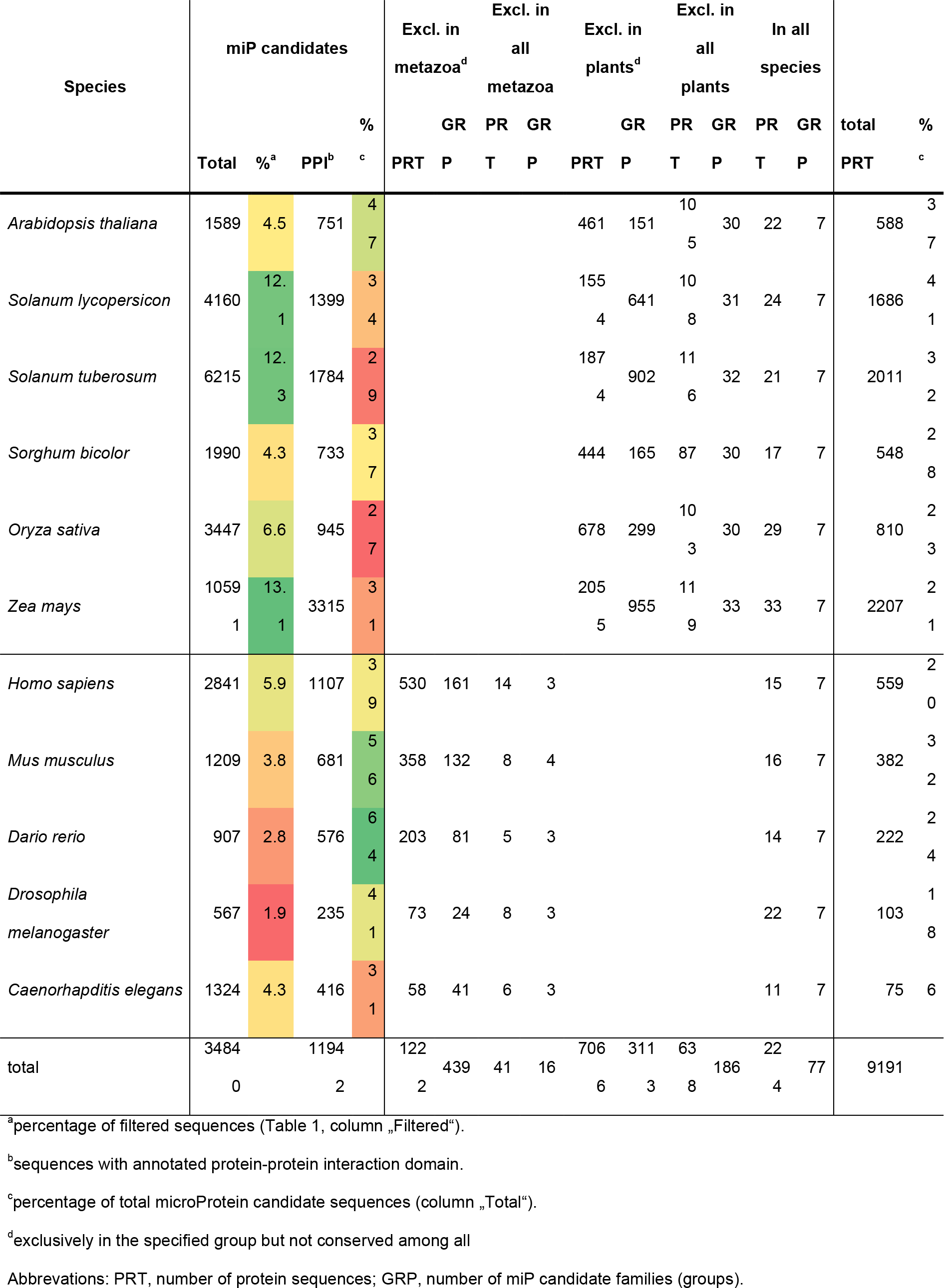
Conserved microProtein candidates.

**Table 3.**
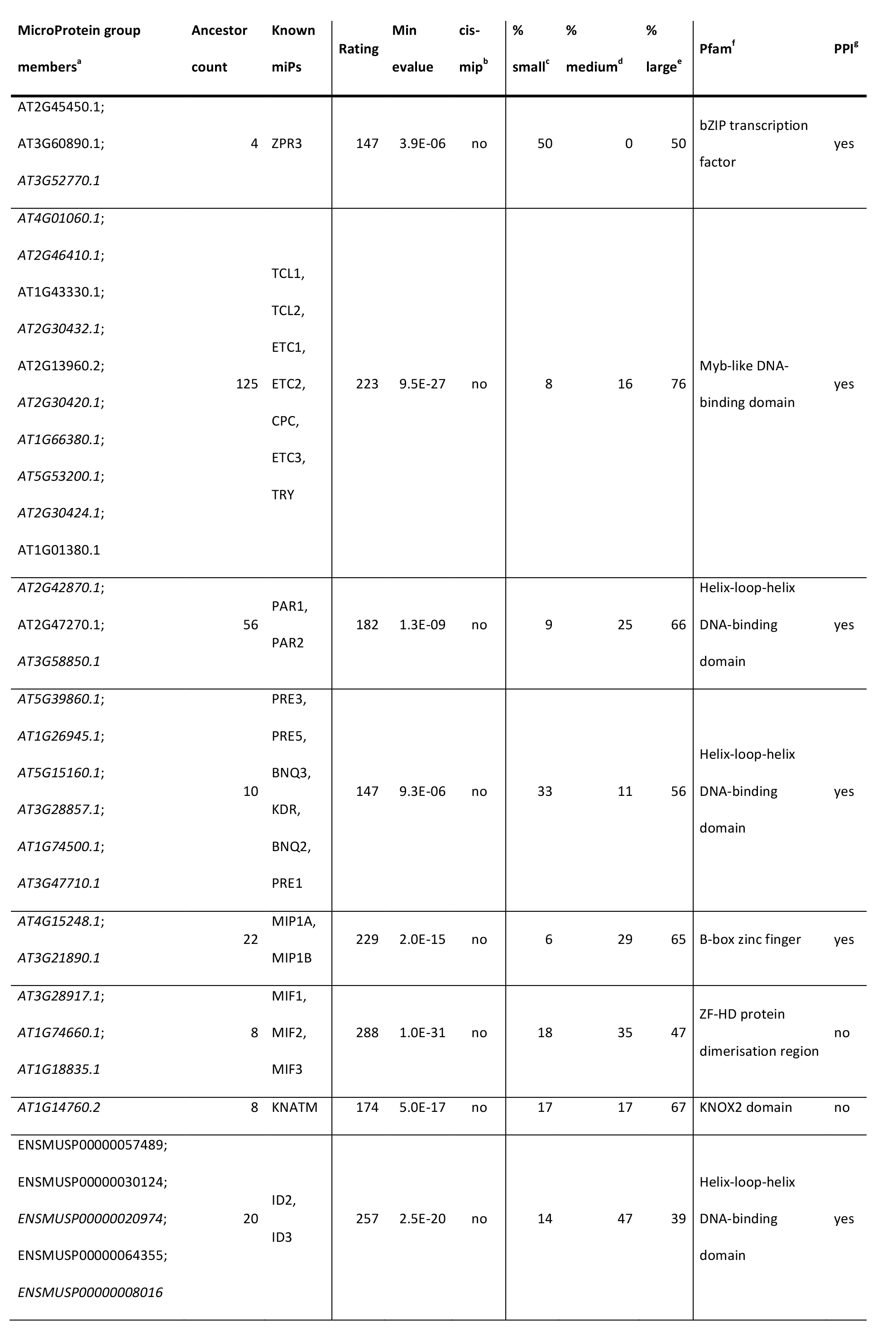
Known microProteins identified by miPFinder.

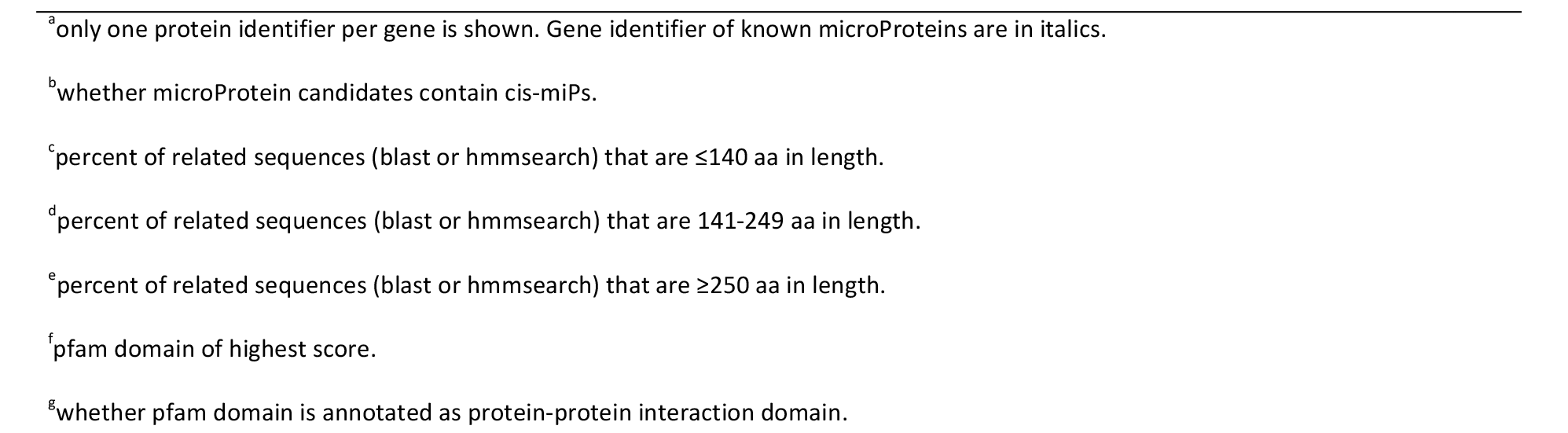

Furthermore, we also identified microProteins that are exclusively found either in plants or metazoans. Each plant dataset contains several hundred to thousands of microProtein candidates that are exclusively conserved among plants. Around 100 proteins in approx. 30 microProtein candidate families per plant have related sequences in all other plants (Fig 2, dark green; Table 2). One third of these are DNA-binding or transcription factor related domains, such as MYB, helix-loop-helix, or zinc finger. Much more microProtein candidates, ranging from 444 in *Sorghum bicolor* to 2,055 in maize, are conserved in at least two plant species (Fig 2, light green; Table 2).

In metazoans, only very few, around 10 microProtein candidates, are conserved in all analyzed genomes (Fig 2, dark blue; Table 2). These sequences have similarity to three structures, the nuclear transport factor 2 (NTF2) domain, ankyrin repeats, and the PDZ domain. The NTF2 families consist in majority of small proteins in contrast to the other two families, which have less than one tenth of small protein sequences. Some dozen to hundreds (from *C. elegans* with 58 to 530 in human) of proteins are conserved exclusively among at least two of the five metazoan proteomes (Fig 2, light blue; Table 2). These numbers differ considerably from microProteins in plants, which might be caused by a bigger evolutionary distance between the chosen metazoan genomes than between the relative closely related plant genomes.

Metazoan microProtein candidates and their putative ancestors were classified into six transcription factor groups and 70 families according to AnimalTFDB (Zhang et al. 2015). Around 10% of microProtein candidate families (Human 117, mouse 54, zebrafish 40, *D. melanogaster* 17, *C. elegans* 43) contained at least one transcription factor (TF). TF Basic Domains Group (e.g. bZIP), Helix-turn-helix (e.g. MYB, homeobox), Other Alpha-Helix Group and Zinc-Coordinating Group (e.g. zf-C2H2) have microProtein candidates in all investigated metazoans (Fig. 3). Some TF families with microProtein candidates were present in several species (e.g. SAND, DM, bHLH, zf-GATA) and few were species specific (e,g, SRF and RFX in human, E2F in mouse, NF-YA in *C. elegans)* (Fig. 3).

**Figure 3.**
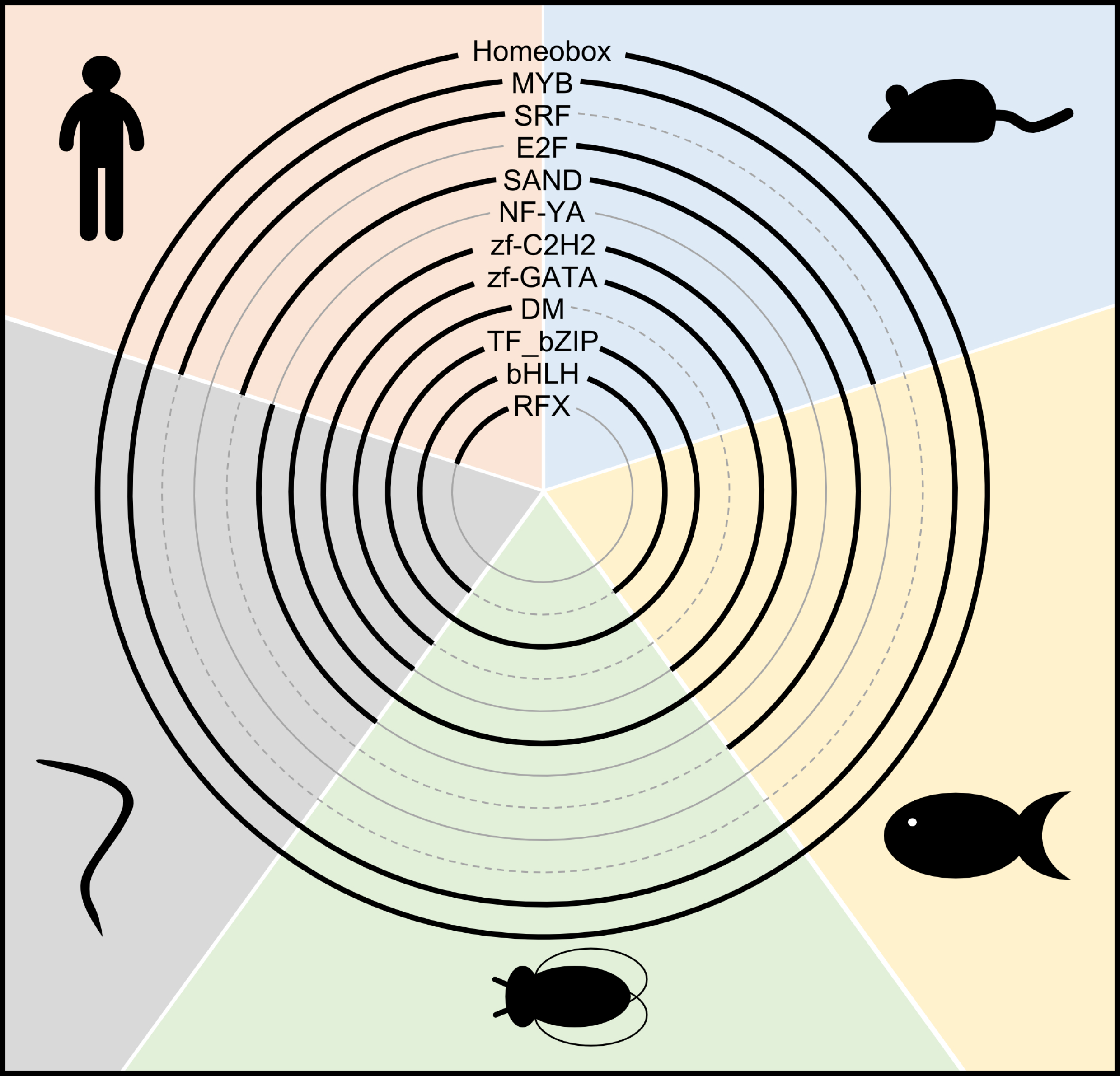
MicroProtein candidates in transcription factor families in metazoans. The presence of microProtein candidates in human (upper left, red), mouse (upper right, blue), zebrafish (right, yellow), fruit fly (bottom, green) and roundworm (left, grey) in the respective transcription factor family is indicated as bold line.

Gene Ontology (GO) terms describe gene products in terms of their associated biological processes, cellular components and molecular functions in a species-independent manner. MicroProtein candidate families were divided into several subsets based on their conservation according to OrthoFinder. For all sets, only the most significant ancestor of each microProtein candidate family was analyzed (Fig. 4). According to GO classifications, many microProtein ancestors are located in the nucleus throughout the subsets, are involved in DNA binding and in protein complexes. The high abundance of proteins with transcription factor properties is in line with the fact that all known microProteins target transcription factors. The biological process 'Anatomical Structure Development' is mostly annotated for metazoan proteins, but also present in dicots but not in monocots. In plant and some metazoan subsets many proteins are involved in response to stress (Fig. 4B). These results support the function of the ancestral genes of known microProteins which are involved in signal transduction, stress responses, and development.

**Figure 4.**
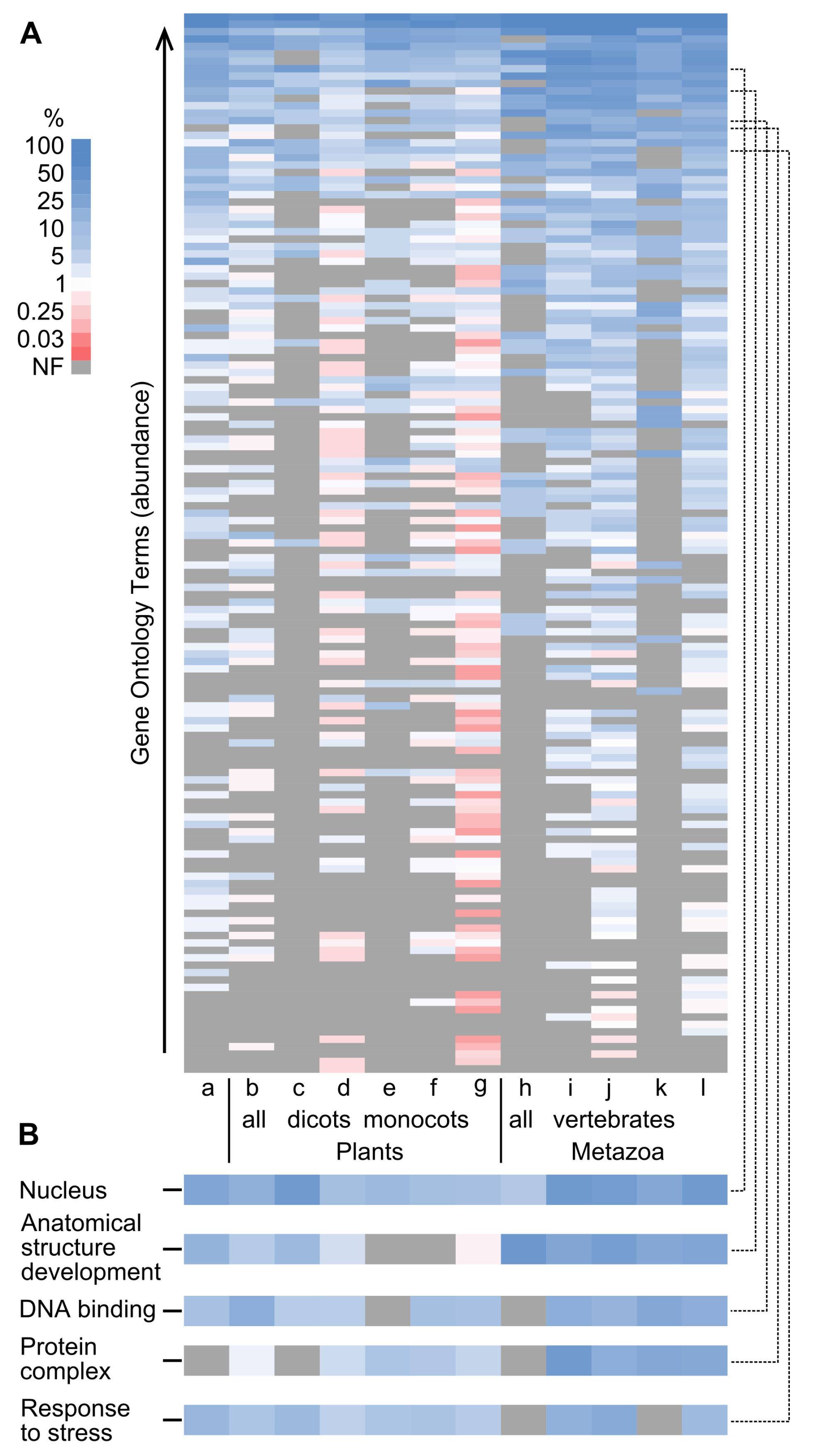
Gene Ontology analysis of microProtein-subsets. For all sets, only the most significant ancestor of a microProtein candidate family was analyzed. The subsets represent microProtein candidate families with the following conservation in: a: all species; b: all plants; c: all dicots; d: some dicots; e: all monocots; f: some monocots; g: some plants; h: all metazoa; i: all vertebrates; j: some vertebrates; k: non-vertebrates; l: some metazoa. A: The GO terms are sorted descending by their average abundance of all subsets and color coded by their subset specific percent of genes with GO annotation. B: Selected GO terms extracted from A as indicated by dashed lines. NF: not found.

## DISCUSSION

We have developed miPFinder, a program to identify microProteins, which are novel regulators of protein function. MiPFinder starts with a set of protein sequences and considers information about protein size, sequence similarity and domain composition to create a list of microProtein candidates. Additionally, when combined with protein conservation information, miPFinder can discriminate between microProteins that occur in several species or microProteins that are species-specific. This resource will aid the identification of microProteins and will promote research on the function of novel microProteins.

An earlier version of miPFinder identified the Arabidopsis microProteins MIP1A and MIP1B (Graeff et al. 2016) that control flowering by recruiting a known flowering activator into a repressive complex. This earlier version was used to identify transcription factor-related microProteins. This new version extends the microProtein concept to any protein harboring a protein-protein-interaction domain. The algorithm is freely available on our webpage (http://cpsc.ku.dk/meet-the-scientists-page/stephan-wenkel-group/miPFinder_v1.zip) and can be used without any restrictions.

Using miPFinder, we screened selected metazoan and plant genomes for microProteins and found that all 22 known Arabidopsis microProteins were identified. Two known metazoan microProteins, mouse ID2 and ID3, are correctly reported but other known microProteins do not fit miPFinder’s size requirements, for example mouse microProtein ID1 which is too large (148aa) or the human RGS5 ancestor protein which is too small (201aa) to fit our parameters. However, the standard settings regarding protein size that we have chosen for this analysis can be easily modified using the appropriate command line parameters in miPFinder.

Several known Arabidopsis microProteins can be found in either all of the six plant genomes that we have investigated here or all at least in one of the sub-sets of the three dicot or monocot genomes. MicroProtein candidates that are conserved among all investigated species seem less likely to have microProtein function because related sequences of these proteins are overall relatively small and larger protein sequences are only distantly similar. In general, we find that microProteins that are conserved in at least a few other species have an increased probability that the small, often one intron sized microProtein candidates are not pseudo-genes. Consequently, microProtein candidate families that are conserved in several but not all of the 11 genomes are promising candidates. Good examples are the LITTLE ZIPPER microProteins, which regulate leaf development and that are conserved in the whole plant euphyllophyte clade. MIP1A/B, which have been shown to fine-tune flowering of Arabidopsis are conserved in all dicotyledonous plants.

We categorized the most significant ancestor of each microProtein candidate family into functional groups and performed a Gene Ontology (GO) analysis. MiPFinder results showed high percentage of GO terms that are also found among ancestors of known microProteins such as 'signal transduction' and 'anatomical structure development' and several 13 microProtein candidates that are related to transcription factors. This pinpoints the importance of microProteins in response to the environment and basic patterning pathways which is exemplified in the role of known microProteins, such as ZPR in Arabidopsis leaf development.

Since known plant microProteins are involved in regulation of transcription, we compared our miPFinder results for metazoa to a transcription factor database. Putative microProteins are present in several major transcription factor families in all studied metazoan genomes. However, in total only 10% of the identified microProtein candidate families are related to transcription factors. This further supports our hypothesis that regulation of protein activity by microProteins extends beyond the regulation of transcription factors and might affect other biological pathways.

The identification and experimental characterization of novel microProteins, based on miPFinder, will allow further improvement of the program. Knowledge of more microProteins will aid in refining the parameters in order to further concentrate the list of microProtein candidates. Additionally, future upgrades of the source databases will benefit microProtein identification. Most importantly a complete and accurate annotation of all small transcripts and respective protein sequences including splice variants will allow for better microProtein detection.

Because microProteins act by engaging in direct protein-protein interactions, candidates with similarity to a known protein-interaction domain are more promising than those without any known domains. MiPFinder annotates protein domains to a given set of sequences, but already existing domain information can also be provided if desired. However, some proteins interact via discontinuous sequences that form three-dimensional interaction interphases rather than with specific interaction domains. These interactions are currently not addressed because interaction networks are not yet completely uncovered and interaction based on structural information is not available for all protein sequences. Due to these constraints, miPFinder does not filter for interaction abilities, it simply annotates potential common interaction domains of microProteins and their related large sequence. Thus highest priority can be given to microProtein candidates with known interaction domains but the search also includes all other candidates.

In summary, selecting microProteins from miPFinder for experimental validation is ideally guided by taking all the above-mentioned criteria into account. For example, MIP1A and MIP1B related protein sequences are in majority large in size (65%), the relative fraction of microProteins is small (6%), and the sequence similarity is rated high. Additionally, MIP1A and MIP1B resemble an annotated protein-protein-interaction domain (B-box) and are exclusively conserved among all three investigated dicots (Arabidopsis, tomato and potato). Therefore these candidates fit perfectly into the scheme of potential microProteins and were experimentally confirmed to have microProtein function. However, when searching for microProteins with a specific function or protein category other priorities might be applicable. Taken together, miPFinder allows the rapid identification of novel microProtein regulators and can be applied to any close-to-complete genome. All settings are adjustable thus allowing users to perform a variety of searches according to their needs. Up to date, microProteins are under-investigated in animals compared to plants and miPFinder enables the identification of microProteins in all available genomes.

## METHODS

### miPFinder script and required standalone applications

The program is written in python v2.7.9 (Python Software Foundation. Python Language Reference, version 2.7. Available at http://www.python.org) and tested for Windows 7. MiPFinder requires the standalone applications hmmer3 (http://hmmer.org/), clustalw2 (http://www.clustal.org/clustal2/) (Larkin et al. 2007) and blast2+ obtained from NCBI (ftp://ftp.ncbi.nih.gov/) (Camacho et al. 2009). These applications are required and have to be installed separately. Sequence files and databases are not provided, the versions used for the analysis herein are described below. The miPFinder script does not include the filter for full-length mRNA sequences, because the optimal procedure differs between organisms and sequence sources, however, a separate script is available.

### Database dependencies

The interaction domain database iPfam v1.0, June 2013, was obtained from http://www.ipfam.org/ (Finn et al. 2014) and Pfam-A_v28.hmm downloaded from Pfam’s FTP site (ftp://ftp.ebi.ac.uk/pub/databases/Pfam/) (Finn et al. 2016). Plant sequence files were downloaded from Pythozome v11 (http://www.phytozome.net) (Goodstein et al. 2012) (Athaliana_167_TAIR10, Osativa_323_v7.0, Sbicolor_313_v3.1, Slycopersicum_225_iTAGv2.3, Stuberosum_206_v3.4, Zmays_284_5b+) and metazoa sequence files were obtained from Ensembl Genes 83 using biomart (http://www.ensembl.org/biomart) (Caenorhabditis_elegans.WBcel235, Danio_rerio.GRCz10, Drosophila_melanogaster.BDGP6, Homo_sapiens.GRCh38, Mus_musculus.GRCm38).

### Circos plot

Only microProtein candidates with similarity to annotated interaction domains (iPfam v1.0, June 2013) were chosen. Orthologous protein sequence families were identified by OrthoFinder v0.3.0 (Emms and Kelly 2015). The data is visualized using Circos v0.68 (Krzywinski et al. 2009).

### Gene Ontology analysis

Gene Ontology (GO) terms for metazoan protein databases were obtained from ENSEMBL and Plant GO terms were retrieved from AgriGO v1.2 (Du et al. 2010). Finally, GOSlimViewer with the generic GOSlim Set from AgBase v2.0 (McCarthy et al. 2006) was used.

### Program usage

The miPFinder program takes a single command line in the windows command prompt (e.g. “python miPFinder.py -f proteins.fasta -p ProteinGeneList.tsv -a annotation.tsv”). The minimum input requirement is a simple fasta file with all protein sequences (“-f”), however a file with protein annotations (“-a”) will aid the microProtein selection tremendously. For the addition of protein-protein interaction domain information, a Pfam domain database (“-d”) and a file specifying interaction domains (“-i”) is necessary. Moreover, a file specifying the protein-gene-relationship (“-p”) will allow for *cis*-microProtein detection, for filtering putative ancestors for their longest splice variant, and for the removal of redundant microProtein candidate splice variants. Parameters for the maximal microProtein and minimum ancestor length can be adjusted (“-M” and “-A”, respectively, standard setting: 140 and 250) as well as all cutoff values.

miPFinder is built with Python v2.7.9 running on Microsoft Windows 7 and using hmmer v3.1b1, blast+ v2.2.29, clustalw v2.1, but any python2, hmmer3, blast2, clustalw2 and Microsoft Windows version might be sufficient. Path to the dependencies (hmmer, blast, clustalw) must be specified, if the accessory programs are not set as environment variables, using command line arguments “-H”,”-B”,”-C”, respectively. MiPFinder will check the availability of specified input files and correct function of all dependencies before each run.

## DATA ACCESS

The miPFinder source code is available on our homepage (http://cpsc.ku.dk/meet-the-scientists-page/stephan-wenkel-group/miPFinder_v1.zip).

## ACKNOWLEDGMENTS

The laboratory of Stephan Wenkel acknowledges funding of this work through a grant from the European Research Council (grant no. 336295) and through Copenhagen Plant Science Centre (CPSC), funded by the University of Copenhagen. We thank Moritz Graeff, Tenai Eguen, Ulla Dolde, Vandasue Rodrigues, Shin-Young Hong, and Esther Botterweg Paredes for commenting on the manuscript and valuable discussions on the identification of microProteins.

### AUTHOR CONTRIBUTIONS

DS implemented the program. DS and SW analyzed the data and wrote the manuscript.

### DISCLOSURE DECLARATION

The authors declare that they have no competing interests.

---

**Supplemental Figure 1.** Correlation of the alignment rating to the minimum e-value.

**Supplemental Table 1.** Table of all microProtein candidates.

## REFERENCES

Benezra R, Davis RL, Lockshon D, Turner DL, Weintraub H. 1990. The protein Id: A negative regulator of helix-loop-helix DNA binding proteins. Cell 61(1): 49–59.

Camacho C, Coulouris G, Avagyan V, Ma N, Papadopoulos J, Bealer K, Madden TL. 2009. BLAST+: architecture and applications. BMC bioinformatics 10: 421.

Du Z, Zhou X, Ling Y, Zhang Z, Su Z. 2010. agriGO: a GO analysis toolkit for the agricultural community. Nucleic acids research 38(Web Server issue): W64–70.

Eguen T, Straub D, Graeff M, Wenkel S. 2015. MicroProteins: small size-big impact. Trends in plant science 20(8): 477–482.

Emms DM, Kelly S. 2015. OrthoFinder: solving fundamental biases in whole genome comparisons dramatically improves orthogroup inference accuracy. Genome biology 16: 157.

Finn RD, Coggill P, Eberhardt RY, Eddy SR, Mistry J, Mitchell AL, Potter SC, Punta M, Qureshi M, Sangrador-Vegas A et al. 2016. The Pfam protein families database: towards a more sustainable future. Nucleic acids research 44(D1): D279–285.

Finn RD, Miller BL, Clements J, Bateman A. 2014. iPfam: a database of protein family and domain interactions found in the Protein Data Bank. Nucleic acids research 42(Database issue): D364–373.

Floyd SK, Ryan JG, Conway SJ, Brenner E, Burris KP, Burris JN, Chen T, Edger PP, Graham SW, Leebens-Mack JH et al. 2014. Origin of a novel regulatory module by duplication and degeneration of an ancient plant transcription factor. Molecular phylogenetics and evolution 81: 159–173.

Goodstein DM, Shu S, Howson R, Neupane R, Hayes RD, Fazo J, Mitros T, Dirks W, Hellsten U, Putnam N et al. 2012. Phytozome: a comparative platform for green plant genomics. Nucleic acids research 40(Database issue): D1178–1186.

Graeff M, Straub D, Eguen T, Dolde U, Rodrigues V, Brandt R, Wenkel S. 2016. MicroProtein-Mediated Recruitment of CONSTANS into a TOPLESS Trimeric Complex Represses Flowering in Arabidopsis. PLoS genetics 12(3): e1005959.

Hu W, Ma H. 2006. Characterization of a novel putative zinc finger gene MIF1: involvement in multiple hormonal regulation of Arabidopsis development. The Plant Journal 45(3): 399–422.

Kim E, Magen A, Ast G. 2007. Different levels of alternative splicing among eukaryotes. Nucleic acids research 35(1): 125–131.

Kim Y-S, Kim S-G, Lee M, Lee I, Park H-Y, Seo PJ, Jung J-H, Kwon E-J, Suh SW, Paek K-H et al. 2008. HD-ZIP III Activity Is Modulated by Competitive Inhibitors via a Feedback Loop in Arabidopsis Shoot Apical Meristem Development. The Plant cell 20(4): 920–933.

Krzywinski M, Schein J, Birol I, Connors J, Gascoyne R, Horsman D, Jones SJ, Marra MA. 2009. Circos: an information aesthetic for comparative genomics. Genome research 19(9): 1639–1645.

Larkin MA, Blackshields G, Brown NP, Chenna R, McGettigan PA, McWilliam H, Valentin F, Wallace IM, Wilm A, Lopez R et al. 2007. Clustal W and Clustal X version 2.0. Bioinformatics 23(21): 2947–2948.

Liang Y, Li C, Guzman VM, Chang WW, Evinger AJ, 3rd, Sao D, Woodward DF. 2005. Identification of a novel alternative splicing variant of RGS5 mRNA in human ocular tissues. The FEBS journal 272(3): 791–799.

McCarthy FM, Wang N, Magee GB, Nanduri B, Lawrence ML, Camon EB, Barrell DG, Hill DP, Dolan ME, Williams WP et al. 2006. AgBase: a functional genomics resource for agriculture. BMC genomics 7: 229.

Seo PJ, Hong S-Y, Ryu JY, Jeong E-Y, Kim S-G, Baldwin IT, Park C-M. 2012. Targeted inactivation of transcription factors by overexpression of their truncated forms in plants. The Plant Journal: no-no.

Tominaga-Wada R, Ishida T, Wada T. 2011. New insights into the mechanism of development of Arabidopsis root hairs and trichomes. International review of cell and molecular biology 286: 67–106.

Wang H, Zhu Y, Fujioka S, Asami T, Li J, Li J. 2009. Regulation of Arabidopsis Brassinosteroid Signaling by Atypical Basic Helix-Loop-Helix Proteins. The Plant cell 21(12): 3781–3791.

Wenkel S, Emery J, Hou B-H, Evans MMS, Barton MK. 2007. A Feedback Regulatory Module Formed by LITTLE ZIPPER and HD-ZIPIII Genes. The Plant cell 19(11): 3379–3390.

Wheelan SJ, Marchler-Bauer A, Bryant SH. 2000. Domain size distributions can predict domain boundaries. Bioinformatics 16(7): 613–618.

Zhang HM, Liu T, Liu CJ, Song S, Zhang X, Liu W, Jia H, Xue Y, Guo AY. 2015. AnimalTFDB 2.0: a resource for expression, prediction and functional study of animal transcription factors. Nucleic acids research 43(Database issue): D76–81.

Zhang L-Y, Bai M-Y, Wu J, Zhu J-Y, Wang H, Zhang Z, Wang W, Sun Y, Zhao J, Sun X et al. 2009. Antagonistic HLH/bHLH Transcription Factors Mediate Brassinosteroid Regulation of Cell Elongation and Plant Development in Rice and Arabidopsis. The Plant cell 21(12): 3767–3780.

